# Unraveling the mechanism of 1-deoxynojirimycin (DNJ) accumulation: the role of SWEET3 in mulberry chloroplasts

**DOI:** 10.1101/2025.02.10.637590

**Authors:** Zhen Yang, Shuo Zhao, Yan Li, Yeling Wei, Zhonghao Wang, Xiaoyu Xia, Dong Li, Ningjia He

**Affiliations:** State Key Laboratory of Resource Insects, Southwest University, Chongqing, 400715, China

**Keywords:** mulberry, 1-deoxynojirimycin, transporter, SWEET3, molecular docking

## Abstract

Transporters are crucial for the accumulation and secretion of plant metabolic products. 1-Deoxynojirimycin (DNJ), a natural compound widely distributed in nature and known for its medicinal properties, has production limitations in microorganisms and herbaceous plants. Mulberry leaves, being easy to harvest and growing rapidly, are the main source of DNJ. However, the current production from mulberry is insufficient to meet the demand. Improving DNJ content in mulberry and thereby reducing production costs is a significant challenge. We first used a green fluorescent tag to visualize the localization of DNJ and found that DNJ highly accumulates in the chloroplasts of mulberry leaves but not in callus tissue. The sugar transporter gene SWEET3, which is highly expressed in mulberry leaves but not in callus, was shown to have DNJ transport capability. Molecular docking analysis revealed that the acidic amino acid residues E68 and D126 in the SWEET3 protein are potential binding sites for DNJ, and both neutral and basic mutations at these sites significantly reduced the ability to DNJ-transporting ability of SWEET3. Furthermore, transient overexpression or RNA interference of SWEET3 in mulberry leaves significantly increased or decreased DNJ content, respectively. Similarly, stable overexpression of the SWEET3 in mulberry hairy roots led to an enhancement in DNJ content within the roots. Our results demonstrated that SWEET3 is essential for DNJ accumulation in mulberry. The study represents a breakthrough in increasing DNJ yield in mulberry through the key role of transporters and identify specific molecular targets required for developing mulberry germplasm with increased DNJ levels.

## Introduction

1-Deoxynojirimycin (DNJ) is a naturally occurring alkaloid with the chemical structure 3,4,5-trihydroxy-2-hydroxymethyl-tetrahydropyridine (Supplementary Figure 1A) (Yagi et al., 1976). This structure closely resembles that of glucose (Supplementary Figure 1B), with a nitrogen atom substituted for an oxygen atom in the pyranose ring. DNJ was first isolated and its structure elucidated from the fermentation broth of *Streptomyces* in 1970, since then, it has garnered significant interest as a potential α-glucosidase inhibitor, offering promise for developing blood glucose-lowering medications (Niwa et al., 1970). DNJ is found in various natural sources; beyond microorganisms, it is present in trace amounts in certain herbaceous plants. The concentration of DNJ in the bulbs of *Hyacinthus orientalis* is approximately 0.002% (Asano et al., 1998), in *Adenora tryphilla* var. *japonica*, it is about 0.0009% (Asano et al., 2000), and in *Commelina communis* var. *hortensis*, it ranges from 0.019% to 0.069% (Shibano et al., 2001). The low levels of DNJ produced by microorganisms and herbaceous plants are insufficient for practical production needs. In contrast, the DNJ content in mulberry (*Morus* spp.) is relatively high, ranging from 0.047% to 0.308% in dry weight (Nuengchamnong et al., 2007). Research has shown significant variability in DNJ content among different mulberry varieties (Kimura et al., 2007). Our previous research identified considerable variations in DNJ levels among different tissues within the same variety, DNJ content was the highest in the bud and roots, but it was not detected in the callus (Yang et al., 2023). Additionally, the DNJ content in mulberry leaves fluctuates due to seasonal and temperature changes (Nakanishi et al., 2011). Despite these variations, mulberry leaves remain the primary source of medicinal DNJ, due to their capacity for frequent harvesting and regeneration, which makes them economically viable for commercial production (Parida et al., 2021). Consequently, increasing the concentration of DNJ in mulberry leaves is essential for optimizing the active constituents of medicinal raw materials and enhancing their utilization efficiency (Parida et al., 2021).

Cell membranes act as selective barriers, allowing only small molecules like gases to pass freely, while the movement of most solutes, including ions and large molecules, depends on specific transport proteins embedded within the membrane (Tang et al., 2020). Numerous studies have explored metabolites absorption and transport in plants, delving into aspects such as uptake kinetics, distribution, and accumulation. Such examples include the theanine transporter CsAAP1 (Dong et al., 2019), the neonicotinoid pesticides transporters PIP1;1 and PIP2;1 (Wan et al., 2024), the nicotine transporters NtJAT1 and NtJAT2 (Morita et al., 2009), the cytokinins transporter ABCG14 (Zhao et al., 2023) and PIN2 (Chen et al., 1998), as well as multidrug and toxic compound extrusion (MATE)-type transporters for plant metabolite translocation (Shoji et al., 2009). Recently, a novel family of sugar transporters, designated SWEET (Sugars Will Eventually be Exported Transporters), has been identified in *Arabidopsis* (Chen et al., 2010). This family is involved in various mechanisms of sugar transport essential for plant growth and development (Yuan et al., 2014). For instance, in apples, the sugar transports MdSWEET2e, MdSWEET9b, and MdSWEET15a are implicated in sugar accumulation within the fruit, with MdSWEET9b and MdSWEET15a serving as primary contributors to the phenotypic variation in sugar concentrations across different cultivars (Zhen et al., 2018). In soybeans, the genes GmSWEET10a and GmSWEET10b play a crucial role in transporting sucrose and hexose, influencing sugar allocation from seed coats to embryos and subsequently affecting oil and protein content as well as seed size (Wang et al., 2020). In rice, OsSWEET11 acts as a plasma membrane sucrose transporter (Chen et al., 2012), while OsSWEET14 collaborates with OsSWEET11 to facilitate endosperm development during the grain filling phase (Yang et al., 2018). The precise transport and accumulation of metabolites mediated by these transport proteins are vital for developing economically significant traits in plants. A comprehensive understanding of the underlying DNJ transmembrane transport mechanisms will enhance our ability to efficiently exploit DNJ from mulberry. However, the mechanisms that govern DNJ transmembrane transport in mulberry remain to be fully clarified. Only a recent contribution by Takasu et al. (2020) reported that the SGLT inhibitor and a GLUT inhibitor significantly suppressed the HepG2 cells uptake of DNJ. So far, these studies have not yet been performed in plant.

This study demonstrates that the sugar transporter SWEET3 plays a role in DNJ accumulation in mulberry, enhancing our understanding of the factors that drive DNJ’s differential distribution across mulberry tissues. This finding provides valuable insights into sugar transporter functions, which could support breeding strategies to increase DNJ content in mulberry germplasm and improve economically important forestry traits.

## Results

### DNJ accumulation primarily occurs in the chloroplasts

Previous research has shown that DNJ is primarily synthesized in young leaves (Yang et al., 2023). Nonetheless, investigations into the specific tissues or cells responsible for DNJ storage within mulberry leaves remain unexplored. NBD (7-nitrobenz-2-oxa-1,3-diazol-4-ylcholesterol) is a fluorescent dye used in studying the localization. In this study, it is employed as a fluorescent tag in labeling DNJ, because of its ability to emit green fluorescence when excited by light in the blue range. We assessed the fluorescence intensity of NBD-DNJ across various concentrations to ensure its validity, the linear regression equation and correlation coefficient for the NBD-DNJ standard curve are shown in Supplementary Figure 2. According to previous reports (Zhou et al., 2024), we successfully prepared protoplasts from both mulberry leaf tissues and callus cultures. Upon incubating the callus-derived protoplasts with 100 μM NBD-DNJ for a duration of one hour, no significant green fluorescence was detected in the callus protoplasts (Figure 1A). Similarly, the incubation of mulberry leaf protoplasts with 100 μM NBD monomer for one hour did not result in a notable accumulation of green fluorescent signals (Figure 1B). Conversely, upon incubation with an equivalent concentration of NBD-DNJ, the green fluorescence was observed to co-localize with the intrinsic red autofluorescence of the chloroplasts in mulberry leaf protoplasts, as visualized through fluorescence microscopy (Figure 1C). These findings indicate that DNJ primarily accumulates in the chloroplasts of mulberry leaf cells. Interestingly, it does not accumulate in callus protoplasts, which lack chloroplasts. This observation is consistent with previous studies that reported undetectable levels of DNJ in callus (Yang et al., 2023).

**Figure 1.**
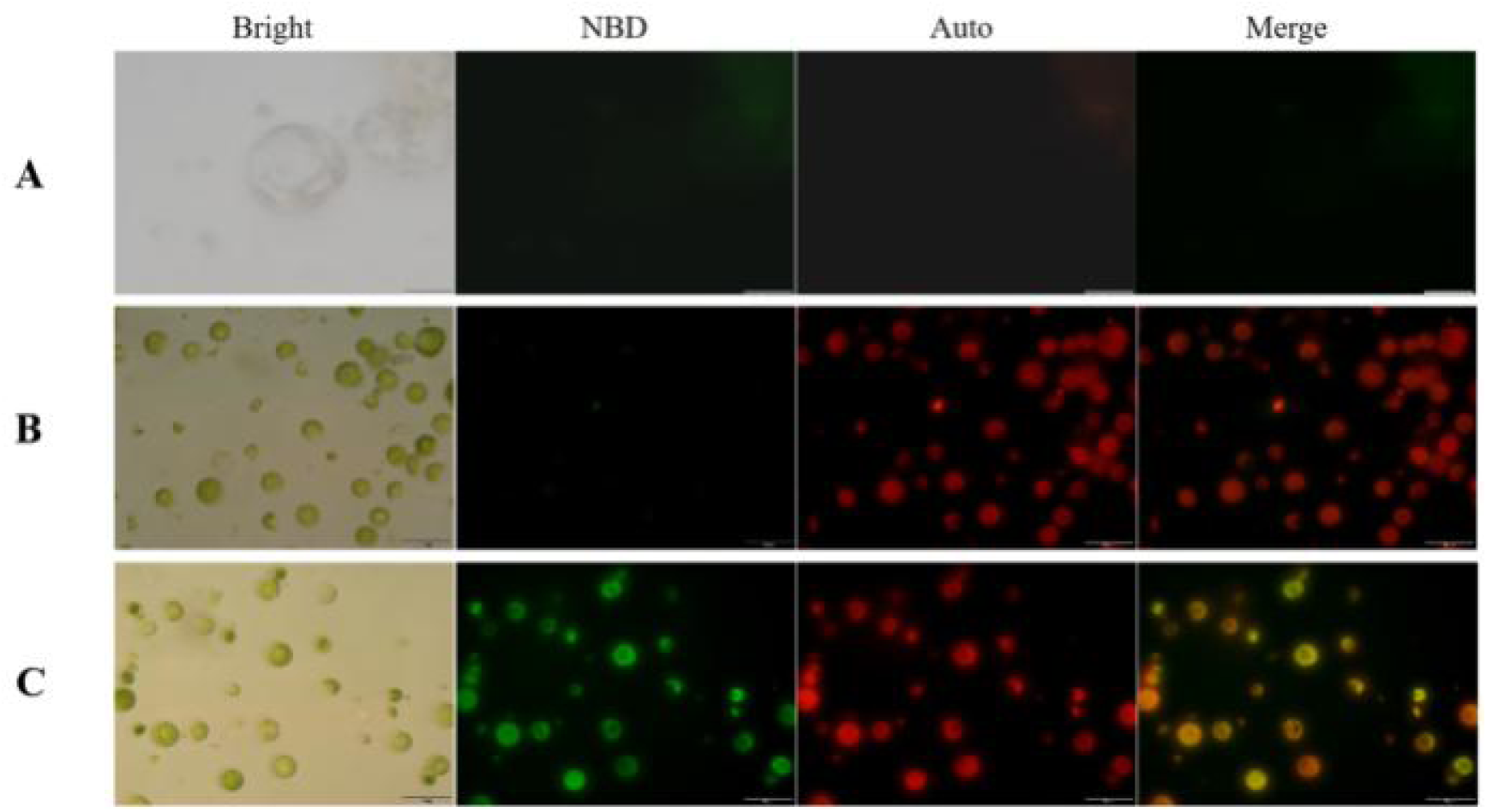
Subcellular localization of DNJ in mulberry protoplast. (**A**) Mulberry callus protoplast incubated with NBD-DNJ (100 µM) alone for 1 hour. Scale bars represent 20 μm. (**B**) Mulberry leaves protoplast incubated with NBD (100 µM) for 1 hour. (**C**) Mulberry leaves protoplast incubated with NBD-DNJ (100 µM) for 1 h. Scale bars represent 50 μm. The confocal images (bright-field and fluorescence at red/green channel) were collected. Auto, autofluorescence image (chloroplasts) in the red channel. NBD (7-Nitrobenz-2-oxa-1,3-diazol-4-yl), a fluorescent probe, were collected at 515-550 nm in the green channel (excitation wavelength: 488 nm). Merged, merged image.

### Expression of sugar transporter genes in mulberry

Based on previous reports suggesting that the accumulation of DNJ in animal cells can be affected by inhibitors of sugar transporters (Takasu et al., 2020), we hypothesize that sugar transporter play a role in the transmembrane transport of DNJ. Considering the conserved nature of ligand-receptor interactions and the substantial structural similarity between DNJ and glucose (Supplementary Figure 1), we hypothesize that members of the sugar transporter family could serve as potential DNJ transporters in mulberry. Based on the observation from Figure 1 that mulberry leaf protoplasts can significantly enrich DNJ while callus-derived protoplasts cannot, we utilized the transcriptome database established from our previous research on DNJ across diverse mulberry tissues (Figure 2A) to identify members of the sugar transporter family that are not expressed in callus tissue but exhibit high expression levels in leaves (Figure 2B). Out of 66 sugar transporters, three transcripts annotated as one putative member of monosaccharide transporter gene family (nucleotide-sugar transporter 6, *NST3*) and two members of the Sugars Will Eventually be Exported Transporters (SWEET) gene family (*SWEET16L* and *SWEET3*), fulfilled the above-mentioned criteria and were selected as candidate genes for further characterization (Figure 3A).

**Figure 2.**
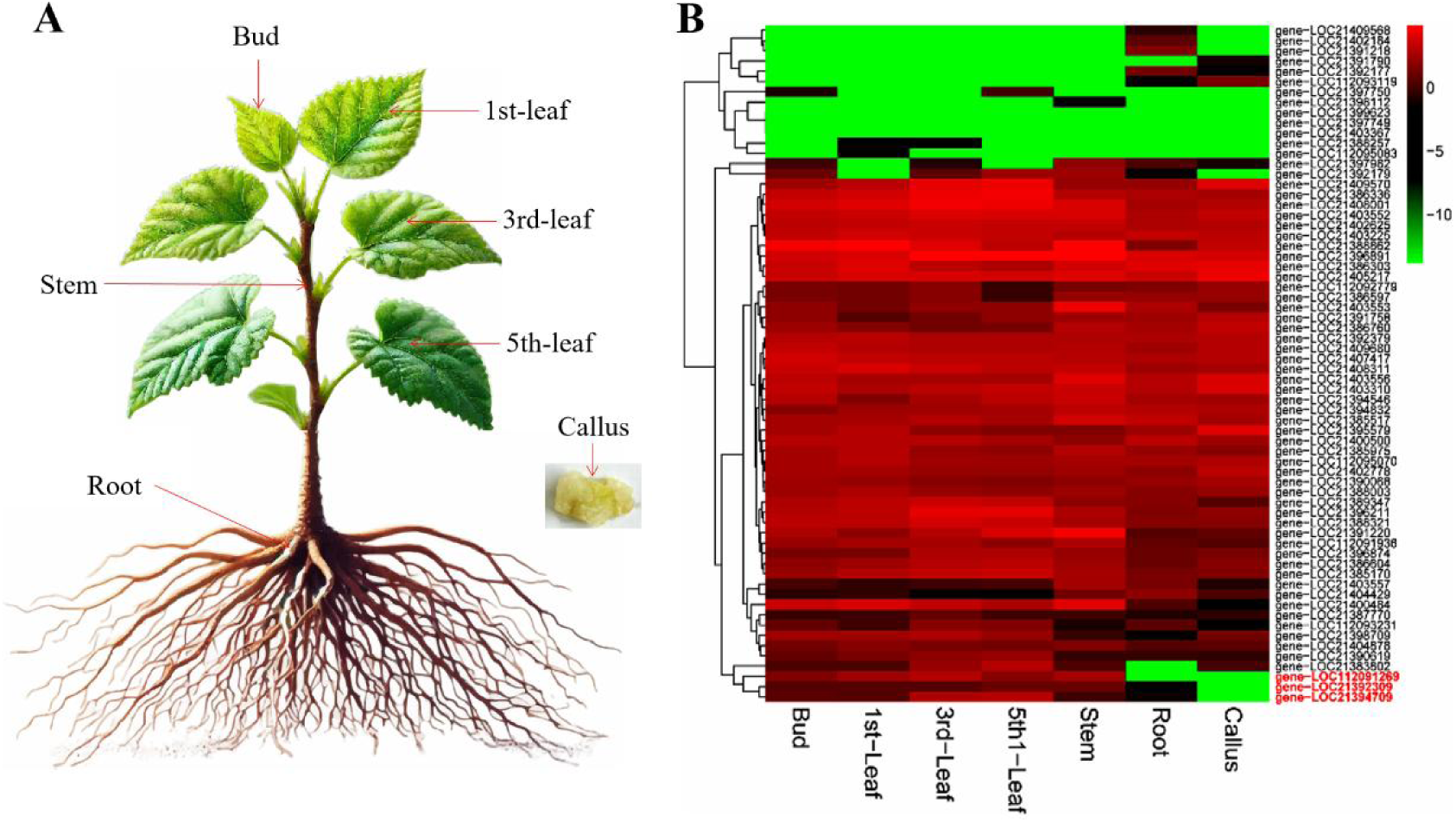
Expression pattern of sugar transporter genes in mulberry. **(A)** Mulberry different tissues. (**B**) Heat map of sugar transporter genes in different tissues of mulberry. The genes highlighted in red are not expressed in callus tissue but show high expression levels in leaves.

**Figure 3.**
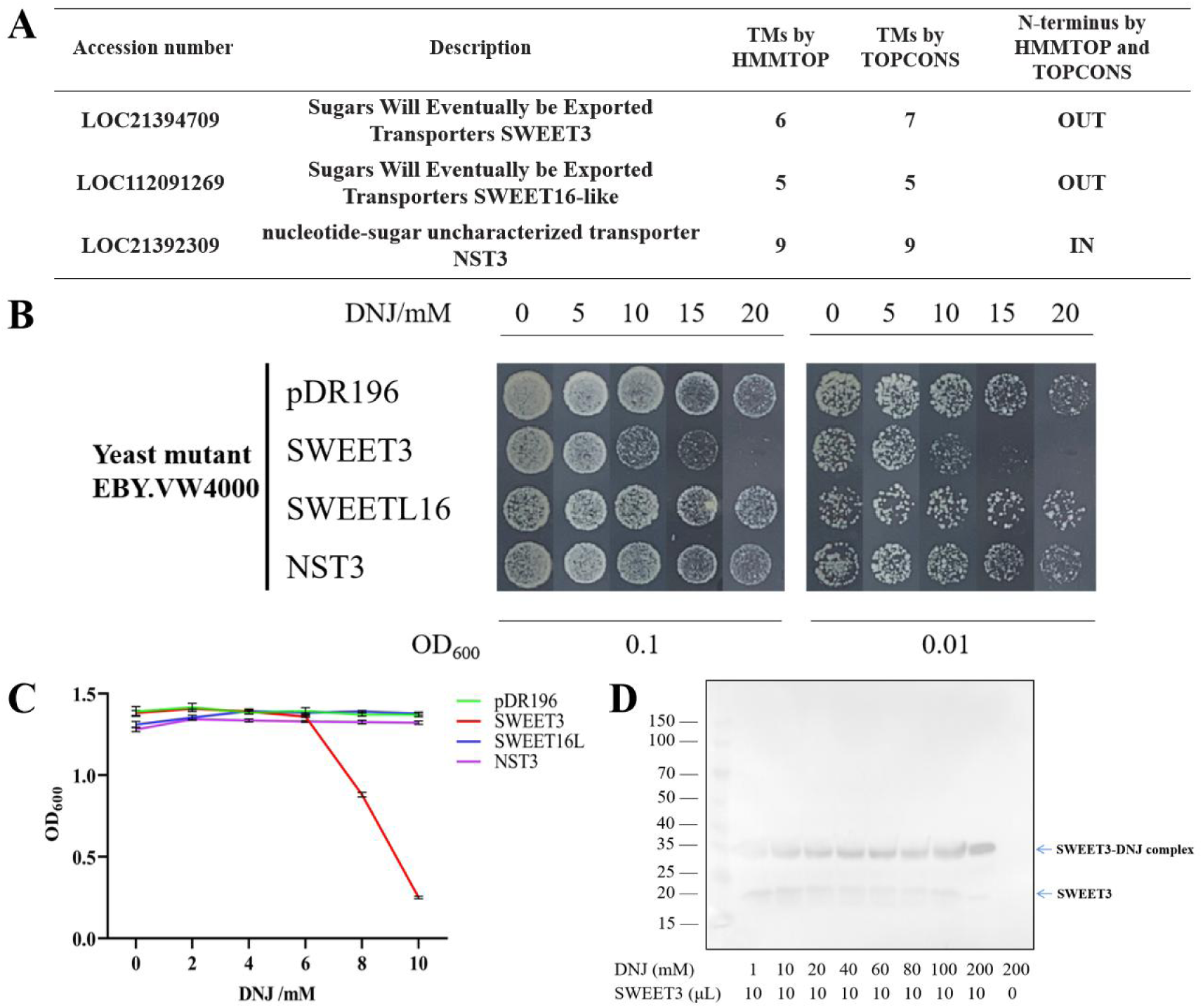
DNJ is specifically transported in yeast by SWEET3. (**A**) Lists of potentially selected candidate sugar transporter genes. The prediction of transmembrane domains (TMs) and N-terminal direction of sugar transporters using two kinds of software HMMTOP and TOPCONS. (**B**) The ability of the tolerance in response to DNJ of *S. cerevisiae* strain EBY.VW4000 expressing SWEET3, SWEET16L and NST3. pDR196, empty vector. (**C**) Yeast growth curve in liquid medium with indicated concentrations of DNJ. Note that 8 and 10 mM DNJ inhibited yeast growth. The OD600 value served as a measure of yeast growth. Data are shown as the mean ± SE (n = 3). (**D**) The gel mobility shifts of the SWEET3-DNJ complex showed a varied range of DNJ.

### SWEET3 functions as a DNJ transporters in *Saccharomyces cerevisiae*

Sugar-transport-deficient *S. cerevisiae* strain EBY.VW4000 has been broadly used as a model to identify Sugar transporters. In our pre-experiment, we established that EBY.VW4000 yeast could not grow on medium with 50 mM DNJ but could grow on medium with 10 mM DNJ (Supplementary Figure 3). Based on this observation, to test for DNJ transport capacity, these candidate genes *NST3*, *SWEET16L*, and *SWEET3* were cloned into the pDR196 vector and then transferred into the yeast EBY.VW4000. Although yeast line EBY.VW4000 could grow on medium with 20 mM DNJ, growth did not occur when it expressed SWEET3 (Figure 3B). In contrast, yeast cells expressing *SWEET16L* and *NST3* exhibit normal growth on 20 mM DNJ, similar to those expressing the empty vector (Figure 3B). We further monitored the growth of yeast strains expressing *NST3*, *SWEET16L* and *SWEET3* in liquid medium with 0, 2, 4, 6, 8, and 10 mM DNJ. The yeast strains were inoculated with an initial OD600 of 0.1 and the growths were measured after cultivation for 24 hours. As shown in Figure 3C, DNJ suppressed the growth of yeast strains expressing *SWEET3* at the low concentrations (8 mM and 10 mM), while the same concentrations did not produce significant effects on growth of EBY.VW4000 expressing *NST3* and *SWEET16L* (Figure 3C). Taken together, these results provided support for the hypothesis that sugar transporter SWEET3 may facilitate the inward transport of DNJ in yeast, and neither of the other two sugar transporters SWEET16L and NST3 exhibited this capability. Furthermore, the DNJ-binding capability of SWEET3 was verified by a gel mobility shift assay. The purified SWEET3 mixed with different concentrations of DNJ was resolved on SDS-PAGE. Compared with the mobility of SWEET3 in the presence of 1 mM DNJ, the binding affinity of SWEET3 significantly increased in a concentration dependent manner by the addition of 10-200 mM DNJ (Figure 3D), which suggested that SWEET3 has the ability to bind DNJ.

Considering recent reports that DNJ metabolism occurs inside bacteria and requires DNJ to be transported across the cell membrane (Zhang et al., 2024), we also focused on the ABC transporter and GABA transporter genes with the same trend in the gene expression of *SWEE3*, that are expressed at low levels in callus tissue but exhibit high expression levels in leaves. A total of four ABC transporter genes (*ABCD2*, *ABCG14*, *ABCG15*, and *ABCG22*) and one GABA transporter gene *GABAT1* were found to be relevant (Supplementary Figure 4A and Supplementary Figure 4B). The recombinant plasmid pDR196-ABCD2, pDR196-ABCG14, pDR196-ABCG15, and pDR196-ABCG22 was introduced into the yeast ABC mutant strain AD1-8 (yor1Δ, snq2Δ, pdr5Δ, pdr10Δ, pdr11Δ, ycf1Δ, pdr3Δ, pdr15Δ), which lacks eight endogenous ABC transporters and is widely used for functional studies of transport proteins. Due to the absence of GABA transport-deficient yeast, the recombinant vector pDR196-GABAT1 was constructed for functional characterization in the AD1-8 yeast strain. Yeast expressing those genes exhibited normal growth, comparable to that of yeast harboring the empty vector pDR196 (Supplementary Figure 4C-D). The results suggest that the ABC and GABA transporters mentioned are not capable of transporting DNJ.

### Identification of key binding sites based on molecular docking analysis

The homology modeling of SWEET3 was performed using an automated protein homology-modeling server SWISS-MODEL (Waterhouse et al., 2018). The molecular structure of DNJ was download from PubChem (Kim et al., 2021). A molecular docking model of SWEET3 with DNJ was performed on CB-Dock2 (Liu et al., 2022) using default parameters. The docking analysis of DNJ with SWEET3 (Supplementary Table 1) revealed that out of five docking poses, two (A and B) exhibited general binding interactions, as indicated by a vina score below the threshold of -5 kcal/mol. As depicted in docking pose A and B, the positively charged imino group of DNJ may interact with two acidic amino acid residues in SWEET3: glutamate at position 68 (Figure 4A and Supplementary Figure 5A) and aspartate at position 126 (Figure 4B and Supplementary Figure 5B). To validate these predicted interactions, site-directed mutagenesis was performed to replace E68 and D126 with neutral (E68Q, glutamine; D126N, asparagine) and basic residues (E68K, lysine; D126H, histidine), as shown in Supplementary Figure 6. The yeast expression vector pDR196 was constructed, and the mutant SWEET3 proteins were expressed in the defective yeast strain EBY.VW4000. The effects of these mutations on DNJ transport were evaluated using spot assays and growth curve analysis. As shown in Figure 4C and 4D, the yeast strains harboring the neutral or basic mutations exhibited increased tolerance to DNJ, suggesting a reduced capacity for DNJ transport. This reduction in transport efficiency was further pronounced in the double mutants. These results indicate that the glutamate residue at position 68 and the aspartate residue at position 126 of SWEET3 are critical for its DNJ transport function.

**Figure 4.**
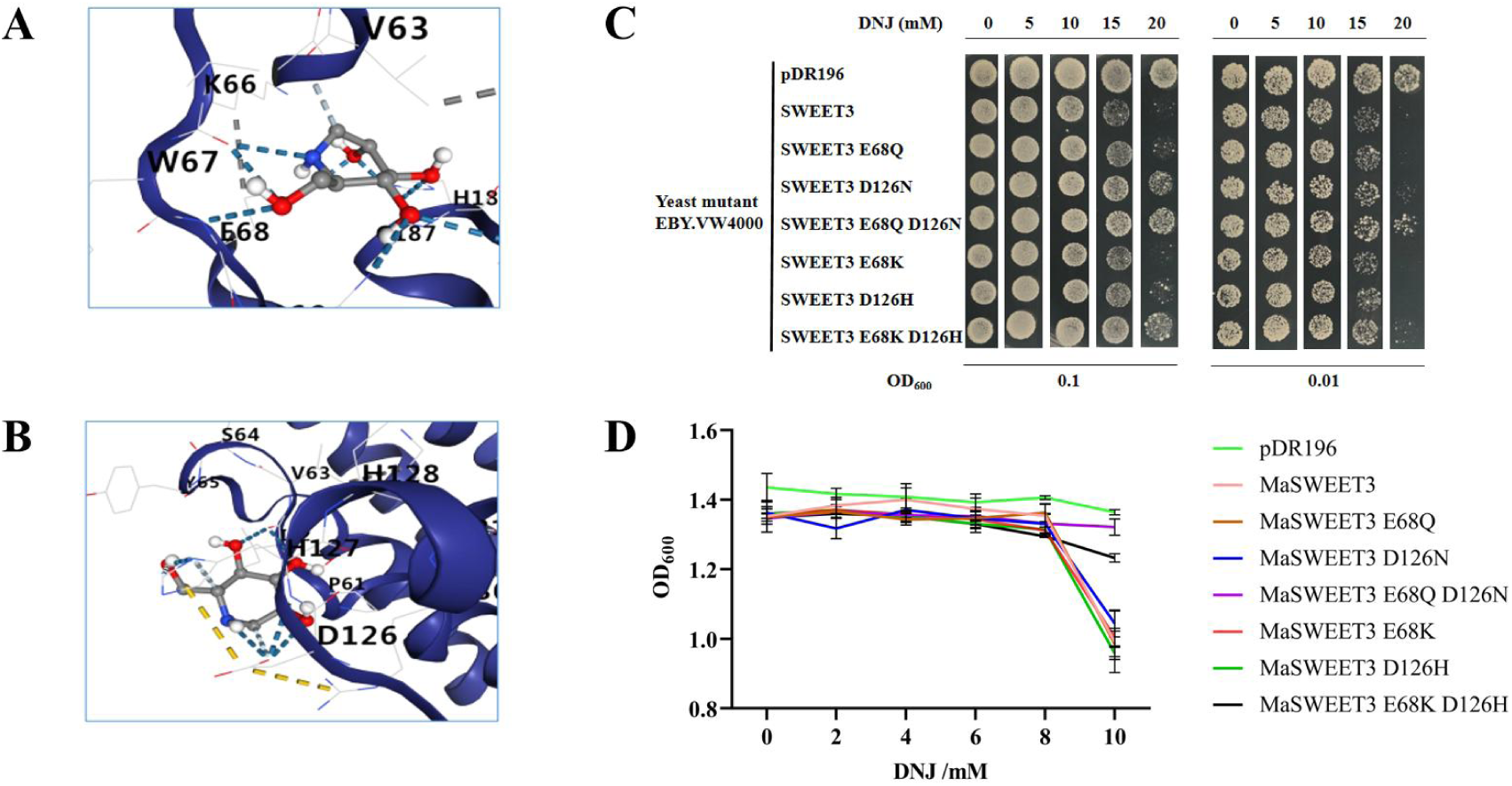
Homology modeling, molecular docking, and site-directed mutagenesis of SWEET3. (**A**) Docking pose A and Docking pose B (**B**) as shown in Table S1. (**C**) Effect of heterologous expression of SWEET3 mutants in EBY.VW4000 on tolerance to DNJ. (**D**) Yeast growth curve for SWEET3 mutants expressed in liquid medium with the indicated concentrations of DNJ. Data are shown as the mean ± SE (n = 3).

### *SWEET3* encodes a chloroplast membrane-localized protein

To explore the structural characteristics of the SWEET3 protein, we utilized the bioinformatics tools HMMTOP (Tusnády and Simon, 2001) and TOPCONS (Tsirigos et al., 2015) to predict its transmembrane domains. The analysis suggested that SWEET3 likely contains 6 or 7 transmembrane regions, with its N-terminus located extracellularly (Figure 3A). Based on this prediction, we examined the subcellular localization of SWEET3 by constructing a fusion protein with GFP, attaching SWEET3 to the N-terminus of green fluorescent protein (GFP) to generate the SWEET3-GFP construct. This construct was transiently expressed in *Nicotiana benthamiana* leaves via *Agrobacterium*-mediated transformation to visualize localization. As shown in Figure 5A, the green fluorescence from the SWEET3-GFP fusion protein co-localized with the red autofluorescence of chlorophyll within chloroplasts, suggesting an association with the chloroplast membrane.

**Figure 5.**
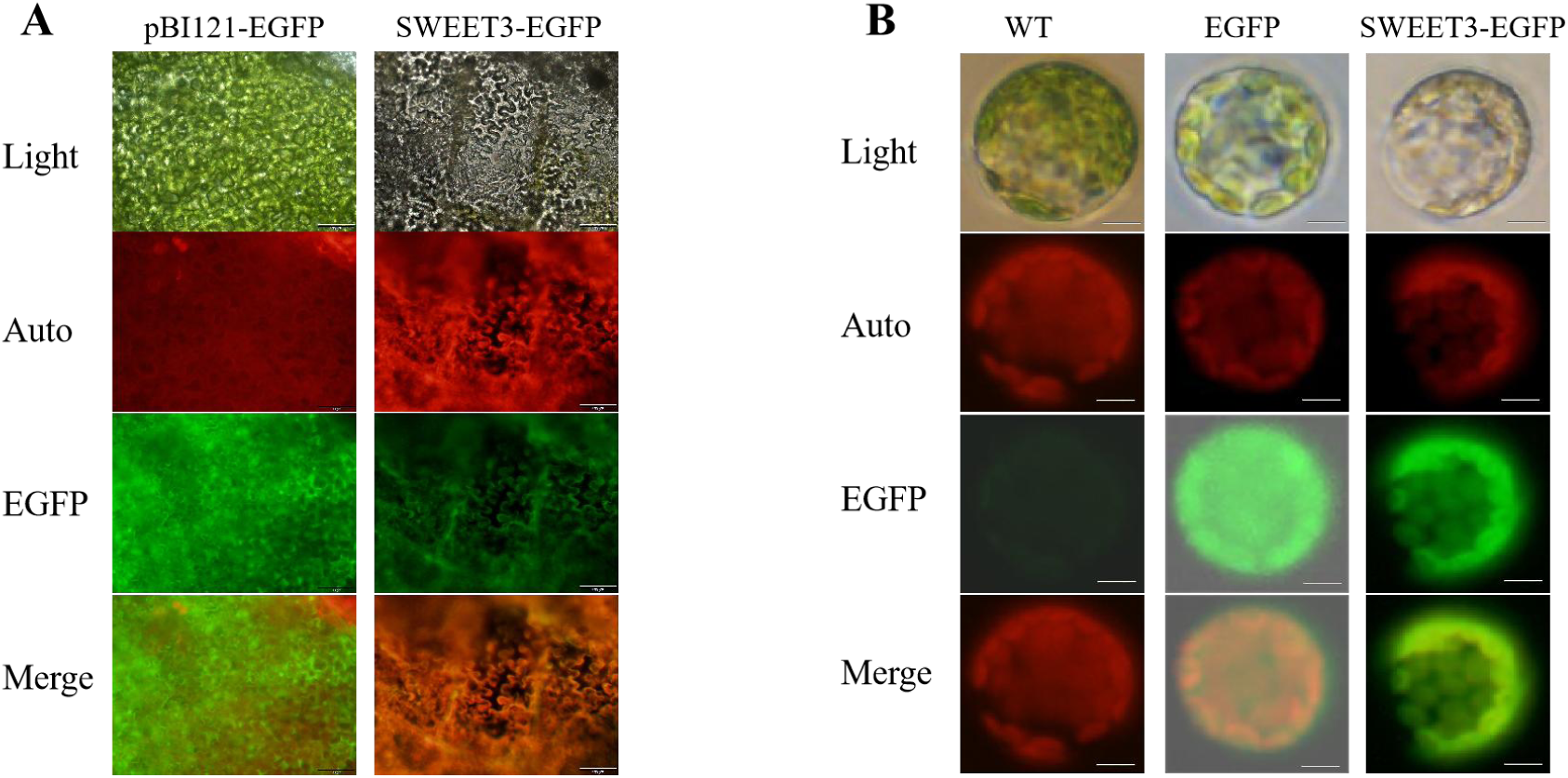
Subcellular localization of SWEET3. (**A**) Confocal micrographs of *Nicotiana benthamiana* leaves transformed with pBI121-EGFP (left) and pBI121-SWEET3-EGFP (right) are shown. Scale bar: 100 μm. (**B**) Confocal micrographs of wild-type (WT) mulberry leaf protoplasts (left), as well as protoplasts transformed with *35S*::GFP (middle) and *35S*::SWEET3-GFP (right), are presented. Scale bar: 10 μm.

To confirm the subcellular localization of SWEET3 in mulberry, we transiently expressed the SWEET3-GFP construct in mulberry leaf protoplasts, following methods outlined in previous studies (Yang et al., 2023; Zhou et al., 2024). As shown in Figure 5B, GFP fluorescence overlapped closely with chlorophyll autofluorescence, supporting the chloroplast membrane localization of SWEET3. These findings suggest that SWEET3 is a membrane-associated protein, specifically localized to the chloroplast membrane, and may play an important role in chloroplast-associated functions.

Given that yeast cells lack chloroplasts, we next sought to determine the subcellular localization of SWEET3 in the yeast mutant strain EBY.VW4000. The pDR196-SWEET3-EGFP vector was constructed for the expression of the SWEET3-EGFP fusion protein, with pDR196-EGFP serving as the control. Both vectors were transformed into the yeast mutant strain EBY.VW4000. Fluorescence signals from the control vector (pDR196-EGFP), containing only the EGFP reporter gene, were observed in the cytoplasm of yeast cells. In contrast, yeast cells expressing pDR196-SWEET3-EGFP exhibited fluorescence signals specifically localized to the plasma membrane (Supplementary Figure 7). This finding regarding localization stands in contrast to the results observed in tobacco and mulberry leaves.

### Significant impact of *SWEET3* expression on DNJ content in mulberry leaves

The functional role of *SWEET3* in mulberry leaves was examined using virus-induced gene silencing (VIGS) and *Agrobacterium*-mediated transient expression techniques (Xin et al., 2021). In the VIGS approach, *SWEET3*-silenced lines were generated, and comparison of these leaves with those injected with the empty vector (2mDNA1) revealed significant reductions in both *SWEET3* expression and DNJ content (Figure 6A-C). To further validate the findings, transient overexpression of *SWEET3* was carried out using an established *Agrobacterium*-mediated expression system. We first construct reporter RUBY that converts tyrosine to vividly red betalain, which is clearly visible to naked eyes without the need of using special equipment or chemical treatments (He et al., 2020). We infiltrated mulberry leaves with Agrobacteria that contain *RUBY*-expressing plasmid, introduction of RUBY made it easier to distinguish the transformed leaves from untransformed leaves (Supplementary Figure 8A-B). We chose the *35S* promoter to drive *RUBY*-P2A-*SWEET3* transient overexpression in mulberry leaves (Figure 6D; Supplementary Figure 8C). RT-qPCR analyses demonstrated a significant increase in *SWEET3* expression in transiently overexpressed leaves compared to control leaves injected with the empty vector (Figure 6E), this upregulation was accompanied by a corresponding increase in DNJ content (Figure 6F). These results collectively highlight that *SWEET3* expression levels play a critical role in regulating DNJ content in mulberry leaves.

**Figure 6.**
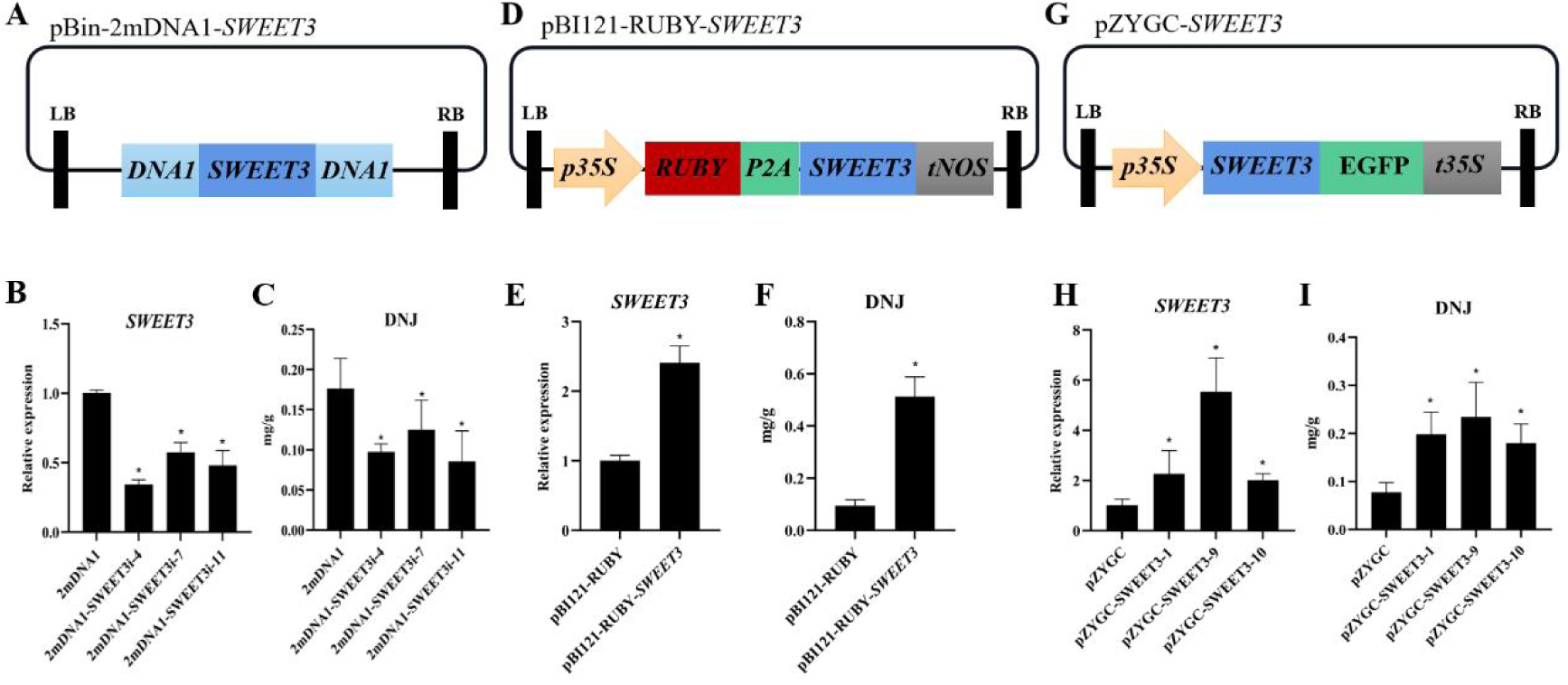
Silencing or overexpression of *SWEET3* affects DNJ accumulation in mulberry. (**A**) Schematic diagram of RNAi vector construction. (**B**) Relative expression of *SWEET3* in RNAi mulberry lines. (**C**) The content of DNJ in RNAi mulberry lines. (**D**) Schematic diagram of transient expression vector construction. P2A, ‘self-cleaving’ 2A peptides. (**E**) Relative expression of *SWEET3* in transient expression mulberry leaves. (**F**) The content of DNJ in mulberry leaves after transient *SWEET3* overexpression. (**G**) Schematic diagram of overexpression vector construction. (**H**) Relative expression of *SWEET3* in overexpression hairy root lines. (**I**) The content of DNJ in overexpression hairy root lines. The metabolite data for each biological line are the mean ± SD of quantification made in three replicates. *p < 0.05 (as determined by a Student’s t-test).

### *SWEET3* overexpression in mulberry roots enhances DNJ accumulation

To further elucidate the function of the *SWEET3* gene, we performed an overexpression study utilizing a hairy root system as a model. We constructed a vector, designated pZYGC, which fused the SWEET3 with enhanced green fluorescent protein (EGFP) to facilitate visualization (Figure 6G). This vector was subsequently transformed into *Agrobacterium rhizogenes* K599 according to established protocols as described previously (Meng et al., 2019; Liu et al., 2021; Yang et al., 2023). After 8 weeks, we successfully obtained transgenic root. Quantitative analysis demonstrated that the relative expression levels of *SWEET3* in the transgenic root lines were significantly higher than those in the control line, as shown in Figure 6H. Importantly, overexpression of *SWEET3* in these hairy roots resulted in DNJ levels that were approximately 1 to 2 times greater than those in wild-type roots, as depicted in Figure 6I. These results collectively indicate that the overexpression of *SWEET3* significantly enhances the accumulation of DNJ in mulberry roots, highlighting its potential role in DNJ transport.

## Discussion

### Sugar transporters are involved in the transmembrane transport of DNJ in both plants and animals

The mulberry is currently recognized as having the highest DNJ content among plants (Zhang et al., 2019; Parida et al., 2021). Mulberry leaves are preferred as the main source of DNJ as they can be harvested more frequently and easily re-grown compared to the other parts of the plant, making them more economically convenient for commercial-scale production (Parida et al., 2021). However, there is a lack of literature addressing the mechanisms of DNJ accumulation in mulberry leaves. Previous studies have shown that DNJ accumulation in leaves increases with higher leaf positions (Nuengchamnong et al., 2007; Vichasilp et al., 2012; Sugiyama et al., 2016). Previous research indicated that DNJ was undetectable in callus tissue induced from leaf explants (Yang et al., 2023), suggesting a possible obstruction to DNJ transport between leaves and the neighboring callus. A previous study demonstrated increased DNJ uptake by colorectal cancer tissues compared to normal tissues in mice (Shuang et al., 2017), it is important to note that cancer tissues exhibit high rates of glucose uptake and elevated expression of sugar transporters to meet their energy demands and support growth (Szablewski, 2013). Preliminary evidence, obtained using sugar transporter inhibitors such as phlorizin and cytochalasins B, suggests that DNJ is mediated by sugar transporters during intestinal absorption and organ uptake in animals (Takasu et al., 2020).

In this study, we identified a gene encoding the Sugars Will Eventually be Exported Transporter SWEET3, which is highly expressed in mulberry leaves abundant in DNJ compared to callus tissues, where DNJ was undetectable. Notably, these callus tissues were directly derived from mulberry leaves under in vitro culture conditions, yet they failed to exhibit substantial SWEET3 expression or DNJ accumulation (Figure 2B). This protein effectively transports and accumulates DNJ in both yeast and mulberry (Figure 3 and Figure 6). A recent study indicated that DNJ metabolism occurs within bacteria and requires DNJ to be transported across the cell membrane, facilitated by the major facilitator superfamily (MFS) transporter and/or GABA permease (Zhang et al., 2024). The MFS is one of the two largest families of membrane transporters found on Earth (Pao et al., 1998), and glucose transporters, members of the MFS, are among the most extensively studied transport proteins (Yan, 2015). Humans and plants use proteins from three major superfamilies for sugar translocation: the MFS, the sodium solute symporter family (SSF; found only in animals), and SWEETs (Tao et al., 2015). In mulberry, the MFS member nucleotide-sugar transporter NST3 lacks the ability to transport DNJ (Supplementary Figure 3), whereas sugar transporter SWEET3, which is not a member of the MFS, is capable of transporting DNJ (Figure 3). Additionally, the low expression of GABA transporters does not account for the inability of callus tissue to transport DNJ across membranes. We also examined another large family of membrane transporters, the ABC transporters, and identified members with low expression in callus tissue, which also failed to exhibit significant DNJ transport activity (Supplementary Figure 4). These findings underscore the crucial role of sugar transporters in the transmembrane transport of DNJ in both plants and animals. Overall, understanding the roles of sugar transporters, especially in DNJ transport, is essential for efficiently obtaining plant-derived DNJ and for optimizing the delivery of DNJ as a therapeutic agent.

### Chloroplast-localized SWEET3 promotes DNJ accumulation in mulberry leaves

The plant genome encodes numerous membrane transport proteins that are essential for fundamental physiological processes, such as ion uptake, metabolite transport, and the excretion and compartmentalization of waste products (Tang et al., 2020). SWEET transporters are versatile in their localization within cellular compartments, primarily found in the plasma membrane (Seo et al., 2011; Kryvoruchko et al., 2016), but also present in the tonoplast (Chardon et al., 2013; Klemens et al., 2013; Guo et al., 2014; Chen et al., 2015) and Golgi membrane (Lin et al., 2014; Chen et al., 2015). However, reports of SWEET transporter localization in chloroplasts remain limited.

In our previous study, we identified the chloroplast as the subcellular localization site for *MnGutB1*, a key gene involved in DNJ biosynthesis in mulberry leaves. Additionally, we found that *MnGutB1* expression is significantly reduced in mature leaves (Yang et al., 2023). In this study, we observed that fluorescently labeled DNJ in the culture medium crossed the cell membrane, entered the protoplasts, and accumulated within the chloroplasts (Figure 1). Moreover, the expression level of the DNJ transporter SWEET3 in the 3rd and 5th leaves was higher than in the young shoots and the 1st leaf (Figure 2B). These findings suggest that DNJ may undergo long-distance transport: after being synthesized in the buds and young leaves, DNJ is transported downward to the lower leaves. This process appears to be mediated by the higher expression of SWEET3 in mature leaves, which supports DNJ content in these leaves and facilitates its accumulation in chloroplasts following transmembrane transport.

Integrating the results from this study with previous findings, we propose a model for DNJ accumulation in mulberry leaf cells. Initially, buds and young leaves synthesize large quantities of DNJ, which then move to lower leaf positions. Newly synthesized DNJ is transported to mature leaves, where SWEET3 is highly expressed in the chloroplasts, mediating DNJ accumulation within the chloroplasts (Figure 7).

**Figure 7.**
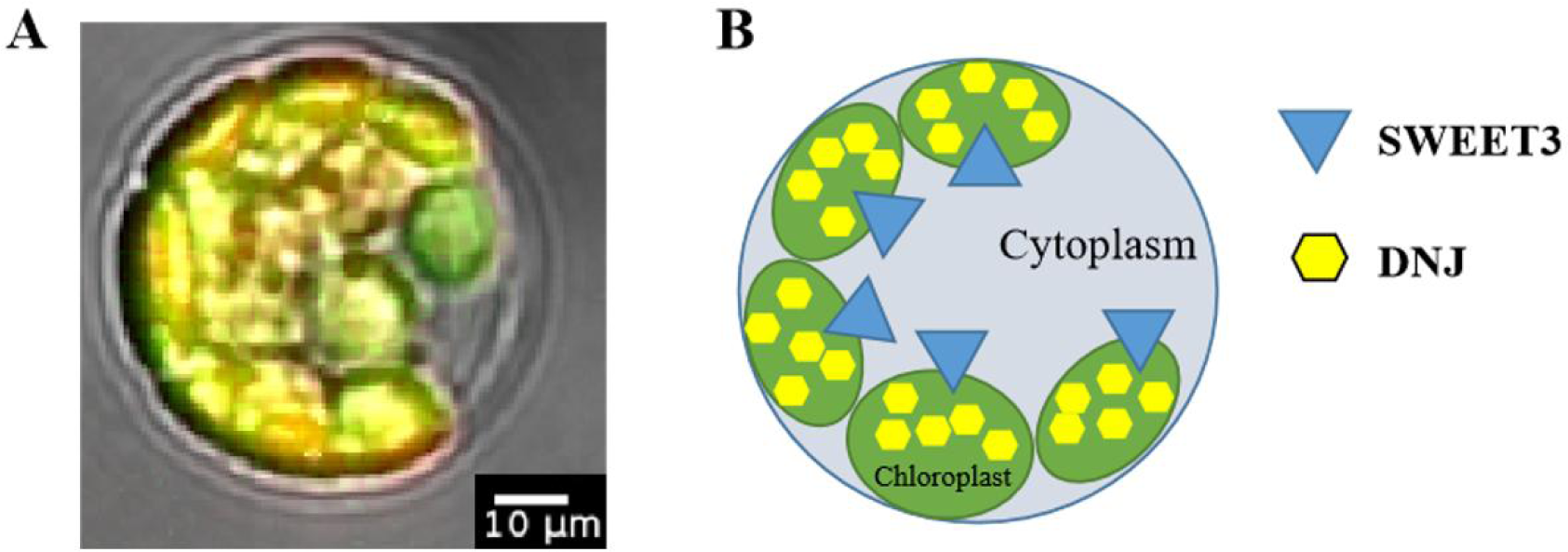
The cellular model of DNJ accumulation in mulberry leaves. (**A**) Colocalization of green fluorescence produced by NBD-DNJ and Chloroplasts autofluorescence. Scale bars represent 10 μm. (**B**) The cellular model of DNJ accumulation mediated by SWEET3 in mulberry leaves.

### Functional analysis of a transporter provides a basis for improving the source of medicinal DNJ

Secondary metabolites in plants are typically produced in low concentrations and are often specific to particular species or confined to narrow taxonomic groups (Gani et al., 2020). Their biosynthesis is frequently compartmentalized within or between cells, necessitating the transport of intermediates and final products through intercellular or intracellular pathways (Tang et al., 2020). The accumulation of these metabolites is tightly regulated by plants via intricate membrane transport systems (Jurgens, 2004; Ayre, 2011; Norambuena and Tejos, 2017; Li et al., 2024). The efficient transport of bioactive compounds is essential for enhancing their utilization efficiency (Morita et al., 2009; Sun et al., 2011; Luo et al., 2020; Zhao et al., 2023; Li et al., 2024; Li et al., 2024). Advanced understanding of these transport systems can provide strategic avenues for the metabolic engineering of medicinal plants, enabling increased production of high-value secondary metabolites (Moses et al., 2013; Yu and De Luca, 2013; Fu et al., 2017; Demurtas et al., 2019).

DNJ is recognized for its hypoglycemic effects and potential therapeutic applications in managing type 2 diabetes mellitus (Liu et al., 2021; Qu et al., 2021). However, the bioavailability and efficacy of DNJ can be influenced by various factors, including the presence of transporters that facilitate its distribution and accumulation in mulberry. By identifying and functionally analyzing these transporters, researchers can develop strategies to improve the delivery of DNJ to target tissues, thereby maximizing its productive potential. The overexpression of the transporter *SWEET3* in mulberry leaves or hairy roots led to a significant increase in DNJ accumulation, exceeding a two-fold enhancement (Figure 6). In our previous study, overexpression of the DNJ biosynthetic gene in hairy roots did not result in such a pronounced increase in DNJ levels (Yang et al., 2023). This suggests that transporter has a more pronounced and direct effect on increasing DNJ content. Furthermore, SWEET3 was expressed on the yeast cell membrane (Supplementary Figure 7), and yeast cells expressing SWEET3 demonstrated the ability to absorb DNJ (Figure 3). This result lays the foundation for the development of engineered strains aimed at the efficient extraction of DNJ.

In conclusion, a comprehensive understanding of transporter mechanisms and their functional analysis is crucial for improving the source of medicinal DNJ. This approach not only paves the way for enhanced therapeutic applications but also contributes to the broader field of drug discovery from natural products.

## Materials and Method

### DNJ visualization and distribution analysis

DNJ standard was sent to Xi’an Qiyue Biotechnology Co., Ltd. for NBD labeling of the green fluorescent group. The labeling efficacy was verified through mass spectrometry and excitation spectra, and a standard curve for NB-DNJ concentration was prepared to assess the labeling ratio of the products. Quality control was performed to ensure standards were met before proceeding with subsequent experimental studies. To observe subcellular localization of the in vitro NBD labeled DNJ in cells, mulberry protoplast cells were incubated with in vitro NBD labeled DNJ (100 μM) or NBD (100 μM) for 1 hour. Fluorescence microscopy was performed with an Olympus IX73 microscope (Olympus, Tokyo, Japan) equipped with an ultrafast three-dimensional fluorescence imaging system (Hooke Instruments, China). The excitation wavelengths and emission filter sets were as follows: GFP, 488 nm (Ex) and 503 to 548 nm (Em); autofluorescence, 644 nm (Ex) and 659 to 700 nm (Em). Image analysis was conducted with the ImageJ software.

### Transcriptome sequencing and analysis

Seven types of tissues were collected from *Morus notabilis*, including young buds, first leaves, third leaves, fifth leaves, fifth leaf veins, stems, roots, and callus induced from in vitro culture, for RNA extraction. The Illumina TruSeq RNA Sample Prep Kit was utilized for library construction. RNA sequencing (RNA-Seq) was performed by Beijing Biomarker Technologies Co., Ltd. The raw sequencing data were filtered to obtain clean data, which were then aligned to the reference genome of *M. notabilis*. The resulting mapped data underwent quality assessments, including insert fragment length checks and randomness tests. Structural analyses were conducted to explore alternative splicing, discover new genes, and optimize gene structures. Differential expression analysis was performed based on gene expression levels across different samples or sample groups, followed by functional annotation and enrichment analysis of differentially expressed genes (DEGs). DEGs were identified based on a false discovery rate (FDR) threshold of <0.05, an absolute log2 fold change value of >1.0, and a p-value of <0.05. Functional annotation was conducted using BLASTx (E-value ⩽ 10^-5) against the NCBI non-redundant protein database, Swiss-Prot protein database, and KEGG database, along with KEGG and Gene Ontology (GO) functional classification. Gene expression levels were calculated using reads per kilobase per million mapped reads (RPKM), and the sugar transporter genes expression matrix was generated for the differential expression analysis (Supplementary Table 2).

### HPLC-ESI-MS/MS hardware parameters and detection conditions

The experimental analysis was conducted using an HPLC-ESI-MS/MS system, specifically the Shim-pack UFLCSHIMADZU CBM20A and Applied Biosystems 4500 QTRAP MS (Chen et al., 2013). A 5 µL sample was injected onto a Waters ACQUITY UPLC HSS T3 C18 column (2.1 mm × 100 mm, 1.8 µm) maintained at 40°C, with a flow rate of 0.4 mL/min. The mobile phase consisted of 0.04% acetic acid in water (Phase A) and 0.04% acetic acid in acetonitrile (Phase B). The elution gradient was set as follows: at 0 min, 95:5 A/B; at 11 min, 5:95 A/B; at 12 min, 5:95 A/B; at 12.1 min, 95:5 A/B; and at 15 min, 95:5 A/B. The eluted compounds were analyzed using ESI-Q TRAP (or QQQ) MS or ESI-Q TOF-MS. The AB Sciex QTRAP 4500 system was equipped with an ESI-Turbo ion source operating in positive ion mode, controlled by Analyst 1.6.1 software (AB Sciex). The operational parameters included an ESI source temperature of 550°C, an ion spray voltage (IS) of 5500 V, curtain gas (CUR) at 25 psi, and collision-activated dissociation (CAD) set to maximum. The QQQ scan was conducted as an MRM experiment, with optimized declustering potential (DP) and collision energy (CE) for each MRM transition. The m/z range was set from 50 to 1000 (Li et al., 2020; Yang et al., 2023).

### Heterologous expression in yeast

The open reading frame (ORF) of transporter genes were amplified by PCR from cDNA in mulberry leaf and cloned into the shuttle vector pDR196 (Fan et al., 2009). PCR primers are listed in Supplementary Table 3. The recombinant vectors or the empty pDR196 vector (control) were separately transferred into the yeast strain EBY.VW4000 or AD1-8 as previously described (Cheng et al., 2015). The transformed cells of the EBY.VW4000 or AD1-8 were grown on synthetically defined uracil dropout medium (SD/-U), and incubated at 30°C for 48 hours. Positive yeast colonies were identified and confirmed via colony PCR and sequencing prior to preservation. The verified yeast strains harboring the pDR196 vector, along with those transformed with the candidate gene, were activated and subsequently diluted to a range of concentrations. These strains were then cultured on SD/-U solid media supplemented with varying concentrations of DNJ (0, 5, 10, 15, 20, and 50 mM). Growth was documented by pictures after 3 days of growth at 30°C. Furthermore, the yeast strains were cultivated in liquid media containing varying concentrations of DNJ (0, 5, 10, 15, and 20 mM) at 30°C for a duration of 24 hours. Subsequently, the optical density at 600 nm (OD600) of the different yeast strains was assessed.

### Molecular docking

Molecular docking was performed to simulate the optimal binding conformation between the protein and the ligand, predicting their binding modes and affinities based on receptor characteristics and interactions with the drug molecule. The structure file of DNJ was obtained from the PubChem database (https://pubchem.ncbi.nlm.nih.gov) (Kim et al., 2021). The complete amino acid sequence of the DNJ transporter was uploaded to the SWISS-MODEL server (https://swissmodel.expasy.org/interactive) for homology modeling (Waterhouse et al., 2018). Molecular docking was then conducted using an online tool CB-Dock2 (http://clab.labshare.cn/cb-dock/php/blinddock.php) (Liu et al., 2022) to simulate the binding conformation of DNJ with its transporter, preliminarily identifying the optimal binding mode and specific binding sites.

### Subcellular localization

Utilizing previously established protocols in our laboratory for the preparation of mulberry leaf protoplasts and PEG-mediated transformation (Zhou et al., 2024), we constructed a recombinant vector containing the candidate transporter gene using the expression vector pAN580 expressing EGFP driven by the *35S* CaMV promoter. Following transformation into mulberry leaf protoplasts and a two-day incubation, the expression localization of the candidate transporter was assessed by observing the position of the green fluorescent protein.

### Electrophoretic mobility shift assay (EMSA) for a metal-binding assay

The metal-binding properties of SWEET3 were confirmed by mixing purified SWEET3 (approximately 0.5 μg/10 μL) with equal volumes of the following solutions: 1, 10, 20, 40, 60, 80, 100 mM DNJ (final concentration). Each mixture was incubated for 30 min at 25 ◦C, was added to 5 μL 5 ×loading buffer, and then was subjected to 10% SDS-PAGE under reducing/nonheating conditions (Ojha and Zhang, 2021).

### VIGS and transient overexpression

Virus-induced gene silencing (VIGS) and transient overexpression was performed following the protocol established by Xin et al. (2021). A 300-bp fragment of the *SWEET3* gene was amplified from *M. notabilis* cDNA via PCR and subsequently cloned into the 2mDNA1 vector, resulting in the plasmid 2mDNA1-*SWEET3*. The Mulberry mosaic dwarf-associated virus (MMDaV) served as the auxiliary plasmid. The plasmids MMDaV, 2mDNA1, and 2mDNA1-*SWEET3* were individually introduced into *Agrobacterium tumefaciens* strain GV3101. A 1:1 (v/v) mixture of MMDaV and 2mDNA1 or its derivatives was infiltrated into the cotyledons of 1-week-old mulberry seedlings using a 1-mL syringe without a needle. After a 24-hour incubation in darkness, the seedlings were transferred to a greenhouse. Thirty days post-infiltration, the second true leaf was collected for analysis using quantitative real-time PCR (qRT-PCR).

RUBY reporter can be used as a visual marker in different crop species (He et al., 2020; Wang et al., 2023). To construct *35s*::RUBY-P2A-SWEET3 plasmid, we linked the RUBY and SWEET3 through 2A peptides into the pBI121 vector between the XhoI and SacI sites. Then, pBI121-RUBY and pBI121-RUBY-P2A-SWEET3 were introduced independently into *A. tumefaciens* strain GV3101, which was then infiltrated into the leaves of 1-month-old mulberry seedlings. The bacterial suspension containing the empty vector pBI121-RUBY was infiltrated into the left side of the leaf, while the recombinant vector pBI121-RUBY-P2A-SWEET3 was infiltrated into the right side. After 24 hours in darkness, the mulberry seedlings were transferred to a greenhouse. Three days post-agroinfiltration, the infiltrated leaves were bisected along the central vein, and the left and right halves were collected separately for analyses of gene expression and DNJ content.

### Transgenic system for mulberry hairy roots

Based on optimized transformation conditions established in our laboratory (Meng et al., 2019; Liu et al., 2021; Yang et al., 2023), the full-length sequence of the transporter gene was cloned into the pZYGC vector and transformed into *Agrobacterium tumefaciens* K599. The bacterial suspension was injected at the junction of true leaves and cotyledons of mulberry plants. After eight weeks of growth, the entire root system of the injected mulberry seedlings was collected. Roots were selected using a fluorescence microscope, and subsequently, they were rapidly frozen in liquid nitrogen for grinding and analysis of transporter gene expression levels and DNJ content.

## Acknowledgments

We greatly appreciate Professor Mohan Gupta (Iowa State University) for sending us the yeast strain AD1-8 and the vector pDR196, Professor Sui Xiaolei (China Agricultural University) for the yeast strain EBY.VW4000.

## Author contributions

Z.Y. and N.H. conceived and designed the research. Z.Y. conducted experiments. Y.W., Y.L., and X.X. provided technical assistance to Z.Y. in the experiments. S.Z., D.L., and Z.W., analyzed the data. Z.Y. wrote the paper, and N.H. revised the manuscript. All authors read and approved the manuscript.

## Data Availability

All experimental data are available and accessible via the main text and/or the supplemental data. The accession number of all sugar transporter genes used in this paper were obtained from the *M. notabilis* Genomics Database (Genome assembly ASM41409v2 https://www.ncbi.nlm.nih.gov/datasets/genome/GCF_000414095.1/) and listed in Supplementary Table 2.

## References

Asano N, et al. (1998) Nitrogen-containing furanose and pyranose analogues from *Hyacinthus orientalis*. J. Nat. Prod. 61: 625–628

Asano N, et al. (2000) Polyhydroxylated pyrrolidine and piperidine alkaloids from *Adenophora triphylla* var. *japonica* (Campanulaceae). Phytochemistry 53: 379–382

Ayre BG (2011) Membrane-transport systems for sucrose in relation to whole-plant carbon partitioning. Mol. Plant 4: 377–394

Chardon F, et al. (2013) Leaf fructose content is controlled by the vacuolar transporter SWEET17 in *Arabidopsis*. Curr. Biol. 23: 697–702

Chen HY, et al. (2015) The *Arabidopsis* vacuolar sugar transporter SWEET2 limits carbon sequestration from roots and restricts *Pythium* infection. Plant J. 83: 1046–1058

Chen L, et al. (2012) Sucrose efflux mediated by SWEET proteins as a key step for phloem transport. Science 6065: 207–211

Chen LQ, et al. (2015) Transport of Sugars. Annu. Rev. Biochem. 84: 865–894

Chen LQ, et al. (2010) Sugar transporters for intercellular exchange and nutrition of pathogens. Nature 468: 527–532

Chen W, et al. (2013) A novel integrated method for large-scale detection, identification, and quantification of widely targeted metabolites: application in the study of rice metabolomics. Mol. Plant 2013: 1769–1780

Cheng JT, et al. (2015) Functional characterization and expression analysis of cucumber (*Cucumis sativus* L.) hexose transporters, involving carbohydrate partitioning and phloem unloading in sink tissues. Plant Sci. 237: 46–56

Demurtas OC, et al. (2019) ABCC transporters mediate the vacuolar accumulation of crocins in saffron stigmas. Plant Cell 31: 2789–2804

Dong C, et al. (2019) Theanine transporters identified in tea plants (*Camellia sinensis* L.). Plant J. 101: 57–70

Fan RC, et al. (2009) Apple sucrose transporter SUT1 and sorbitol transporter SOT6 interact with cytochrome b5 to regulate their affinity for substrate sugars. Plant Physiol. 150: 1880–1901

Fu X, et al. (2017) AaPDR3, a PDR transporter 3, is involved in sesquiterpene β-caryophyllene transport in *Artemisia annua*. Front. Plant Sci. 8: 723

Gani U, et al. (2020) Membrane transporters: the key drivers of transport of secondary metabolites in plants. Plant Cell Rep. 40: 1–18

Guo WJ, et al. (2014) SWEET17, a facilitative transporter, mediates fructose transport across the tonoplast of *Arabidopsis* roots and leaves. Plant Physiol. 164: 777–789

He Y, et al. (2020) A reporter for noninvasively monitoring gene expression and plant transformation. Hortic. Res. 7: 152

Jurgens G (2004) Membrane trafficking in plants. Annu. Rev. Cell Dev. Biol. 20: 481–504

Kim S, et al. (2021) PubChem in 2021: new data content and improved web interfaces. Nucleic Acids Res. 49: 1388–1395

Kimura T, et al. (2007) Food-grade mulberry powder enriched with 1-deoxynojirimycin suppresses the elevation of postprandial blood glucose in humans. J. Agric. Food. Chem. 55: 5869–5874

Klemens PA, et al. (2013) Overexpression of the vacuolar sugar carrier AtSWEET16 modifies germination, growth, and stress tolerance in *Arabidopsis*. Plant Physiol. 163: 1338–1352

Kryvoruchko IS, et al. (2016) MtSWEET11, a nodule-specific sucrose transporter of *Medicago truncatula*. Plant Physiol. 171: 554–565

Li D, et al. (2020) MMHub, a database for the mulberry metabolome. Database 2020: baaa011

Li F, et al. (2024) Characterization of a vacuolar importer of secologanin in *Catharanthus roseus*. Commun. Biol. 7: 939

Li M, et al. (2024) CBL1/CIPK23 phosphorylates tonoplast sugar transporter TST2 to enhance sugar accumulation in sweet orange (*Citrus sinensis*). J. Integr. Plant Biol. 00: 1–18

Li Y, et al. (2024) The ABC transporter SmABCG1 mediates tanshinones export from the peridermic cells of *Salvia miltiorrhiza* root. J. Integr. Plant Biol. 00: 1–15

Lin IW, et al. (2014) Nectar secretion requires sucrose phosphate synthases and the sugar transporter SWEET9. Nature 508: 546–549

Liu CR, et al. (2021) Transcriptome and DNA methylome reveal insights into phytoplasma infection responses in mulberry (*Morus multicaulis* Perr.). Front. Plant Sci. 12: 697702

Liu Q, et al. (2021) Ramulus mori (sangzhi) alkaloids (SZ-A) ameliorate glucose metabolism accompanied by the modulation of gut microbiota and ileal inflammatory damage in type 2 diabetic kkay mice. Front. Pharmacol. 12: 642400

Liu Y, et al. (2022) CB-Dock2: improved protein-ligand blind docking by integrating cavity detection, docking and homologous template fitting. Nucleic Acids Res. 50: 159–164

Luo X, et al. (2020) Antimicrobial peptide reverses ABCB1-mediated chemotherapeutic drug resistance. Front. Pharmacol. 11: 1208

Meng D, et al. (2019) Development of an efficient root transgenic system for pigeon pea and its application to other important economically plants. Plant Biotechnol. J. 17: 1804–1813

Morita M, et al. (2009) Vacuolar transport of nicotine is mediated by a multidrug and toxic compound extrusion (MATE) transporter in *Nicotiana tabacum*. Proc. Natl. Acad. Sci. USA 106: 2447–2452

Moses T, et al. (2013) Bioengineering of plant (tri)terpenoids: from metabolic engineering of plants to synthetic biology in vivo and in vitro. New Phytol. 200: 27–43

Nakanishi H, et al. (2011) Effect of environmental conditions on the α-glucosidase inhibitory activity of mulberry leaves. Biosci. Biotechnol. Biochem. 75: 2293–2296

Niwa T, et al. (1970) “Nojirimycin” as a potent inhibitor of glucosidase. J. Agric. Chem. Soc. Japan 34: 966–968

Norambuena L, Tejos R (2017) Chemical genetic dissection of membrane trafficking. Annu. Rev. Plant Biol. 68: 197–224

Nuengchamnong N, et al. (2007) Quantitative determination of 1-deoxynojirimycin in mulberry leaves using liquid chromatography–tandem mass spectrometry. J. Pharm. Biomed. Anal. 44: 853–858

Ojha A, Zhang W (2021) Characterization of gustatory receptor 7 in the brown planthopper reveals functional versatility. Insect Biochem. Mol. Biol. 132: 103567

Pao SS, et al. (1998) Major facilitator superfamily. Microbiol. Mol. Biol. Rev. 62: 1–34

Parida IS, et al. (2021) A comprehensive review on the production, pharmacokinetics and health benefits of mulberry leaf iminosugars: Main focus on 1-deoxynojirimycin, d-fagomine, and 2-O-α-d-galactopyranosyl-DNJ. Crit. Rev. Food. Sci. Nutr. 63: 3468–3496

Qu L, et al. (2021) Efficacy and safety of mulberry twig alkaloids tablet for the treatment of type 2 diabetes: a multicenter, randomized, double-blind, double-dummy, and parallel controlled clinical trial. Diabetes Care 44: 1324–1333

Seo PJ, et al. (2011) An *Arabidopsis* senescence-associated protein SAG29 regulates cell viability under high salinity. Planta 233: 189–200

Shibano M, et al. (2001) Determination of 1-deoxynojirimycin and 2, 5-dihydroxymethyl 3, 4-dihydroxypyrrolidine contents of *Commelina communis* var. *hortensis* and the antihyperglycemic activity. Natural Medicines 55: 251–254

Shoji T, et al. (2009) Multidrug and toxic compound extrusion-type transporters implicated in vacuolar sequestration of nicotine in tobacco roots. Plant Physiol. 149: 708–718

Shuang E, et al. (2017) Intake of mulberry 1-deoxynojirimycin prevents colorectal cancer in mice. J. Clin. Biochem. Nutr. 61: 47–52

Sugiyama M, et al. (2016) Effect of solar radiation on the functional components of mulberry (*Morus alba* L.) leaves. J. Sci. Food Agric. 96: 3915–3921

Sun A, et al. (2011) A transgenic study on affecting potato tuber yield by expressing the rice sucrose transporter genes OsSUT5Z and OsSUT2M. J. Integr. Plant Biol. 53: 586–595

Szablewski L (2013) Expression of glucose transporters in cancers. Biochim. Biophys. Acta. 1835: 164–169

Takasu S, et al. (2020) Intestinal absorption and tissue distribution of aza-sugars from mulberry leaves and evaluation of their transport by sugar transporters. J. Agric. Food. Chem. 68: 6656–6663

Tang RJ, et al. (2020) Plant membrane transport research in the post-genomic era. Plant Commun. 1: 100013

Tao Y, et al. (2015) Structure of a eukaryotic SWEET transporter in a homotrimeric complex. Nature 527: 259–263

Tsirigos KD, et al. (2015) The TOPCONS web server for consensus prediction of membrane protein topology and signal peptides. Nucleic Acids Res. 43: 401–407

Tusnády GE, Simon I (2001) The HMMTOP transmembrane topology prediction server. Bioinformatics 17: 849–850

Vichasilp C, et al. (2012) Development of high 1-deoxynojirimycin (DNJ) content mulberry tea and use of response surface methodology to optimize tea-making conditions for highest DNJ extraction. LWT - Food Sci. Technol. 45: 226–232

Wan Q, et al. (2024) Two aquaporins, PIP1;1 and PIP2;1, mediate the uptake of neonicotinoid pesticides in plants. Plant Commun. 5: 100830

Wang D, et al. (2023) The RUBY reporter enables efficient haploid identification in maize and tomato. Plant Biotechnol. J. 21: 1707–1715

Wang S, et al. (2020) Simultaneous changes in seed size, oil content and protein content driven by selection of SWEET homologues during soybean domestication. Natl. Sci. Rev. 7: 1776–1786

Waterhouse A, et al. (2018) SWISS-MODEL: homology modelling of protein structures and complexes. Nucleic Acids Res. 46: 296–303

Xin Y, et al. (2021) Dynamic changes in transposable element and gene methylation in mulberry (*Morus notabilis*) in response to Botrytis cinerea. Hortic. Res. 8: 154

Yagi M, et al. (1976) The structure of moranoline, a piperidine alkaloid from *Morus* species. B. Chem. Soc. Jpn. 50: 571–572

Yan N (2015) Structural biology of the major facilitator superfamily transporters. Annu. Rev. Biophys. 44: 257–283

Yang J, et al. (2018) SWEET11 and 15 as key players in seed filling in rice. New Phytol. 218: 604–615

Yang Z, et al. (2023) Dehydrogenase MnGutB1 catalyzes 1-deoxynojirimycin biosynthesis in mulberry. Plant Physiol. 192: 1307–1320

Yu F, De Luca V (2013) ATP-binding cassette transporter controls leaf surface secretion of anticancer drug components in *Catharanthus roseus*. Proc. Natl. Acad. Sci. USA 110: 15830–15835

Yuan M, et al. (2014) Rice MtN3/saliva/SWEET gene family: Evolution, expression profiling, and sugar transport. J. Integr. Plant Biol. 56: 559–570

Zhang N, et al. (2024) Gut bacteria of lepidopteran herbivores facilitate digestion of plant toxins. Proc. Natl. Acad. Sci. USA 121: e2412165121

Zhang W, et al. (2019) An overview of the biological production of 1-deoxynojirimycin: current status and future perspective. Appl. Microbiol. Biotechnol. 103: 9335–9344

Zhao J, et al. (2023) *Arabidopsis* ABCG14 forms a homodimeric transporter for multiple cytokinins and mediates long-distance transport of isopentenyladenine-type cytokinins. Plant Commun. 4: 100468

Zhen Q, et al. (2018) Developing gene-tagged molecular markers for evaluation of genetic association of apple SWEET genes with fruit sugar accumulation. Hortic. Res. 5: 14

Zhou H, et al. (2024) Efficient mesophyll-derived protoplast manipulation system as a versatile tool for characterization of genes responding to multiple stimuli in mulberry. Plant Cell Tissue Organ Cult. 156: 50

