## Supplementary_Figure 1-8 and Table 1-3 for "Unraveling the mechanism of 1-deoxynojirimycin (DNJ) accumulation: the role of SWEET3 in mulberry chloroplasts": Supplementary_Figure.docx

**Supplemental Figure**

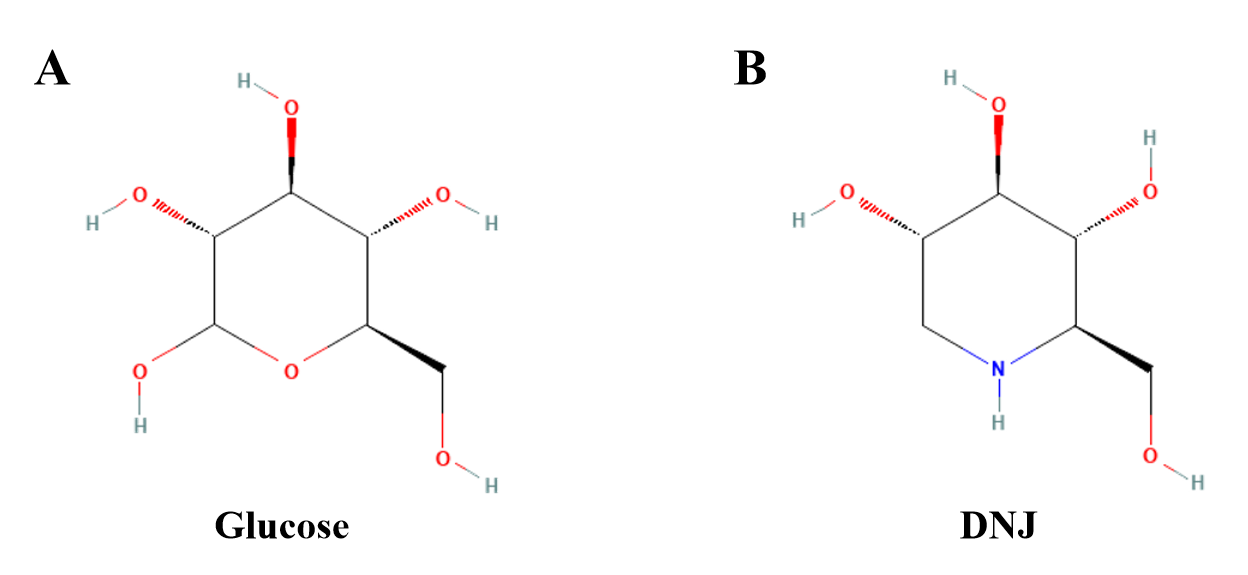

**Figure S1.** **Molecular structural formula of the (A) glucose and (B) DNJ.**

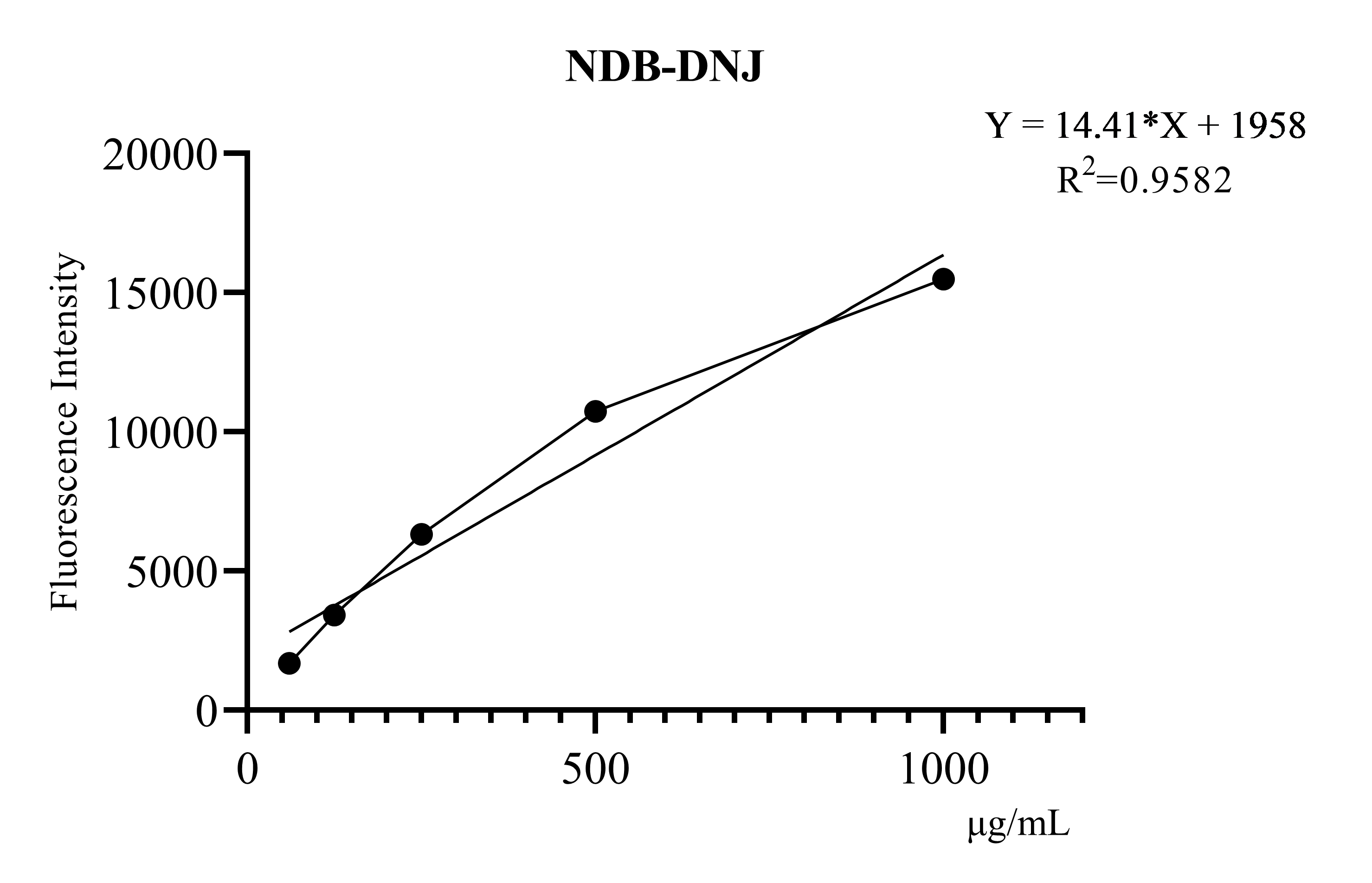

**Figure S2.** **The relationship between the fluorescence intensity and the concentration of NBD-DNJ in the range of 0-1000 µg/mL.**

The linear regression equation and correlation coefficient for the standard curve are shown in in the upper right corner.

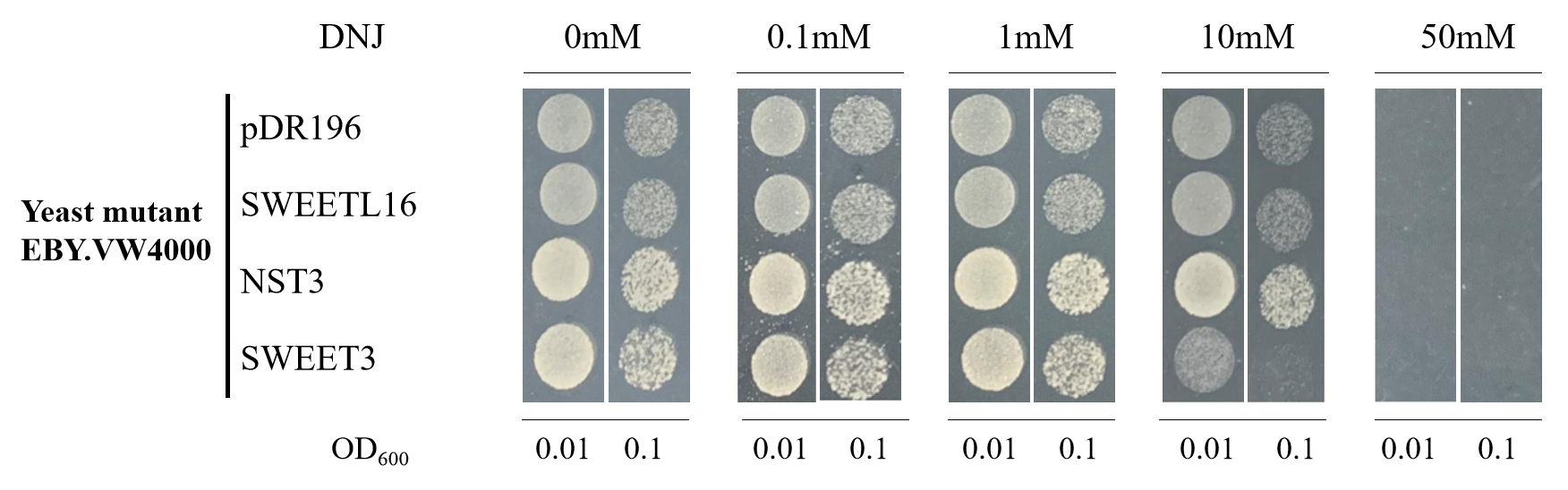

**Figure S3. The tolerance of *S. cerevisiae* strain EBY.VW4000, expressing SWEET3, SWEET16L, and NST3, was evaluated under varying DNJ concentrations ranging from 0 to 50 mM.**

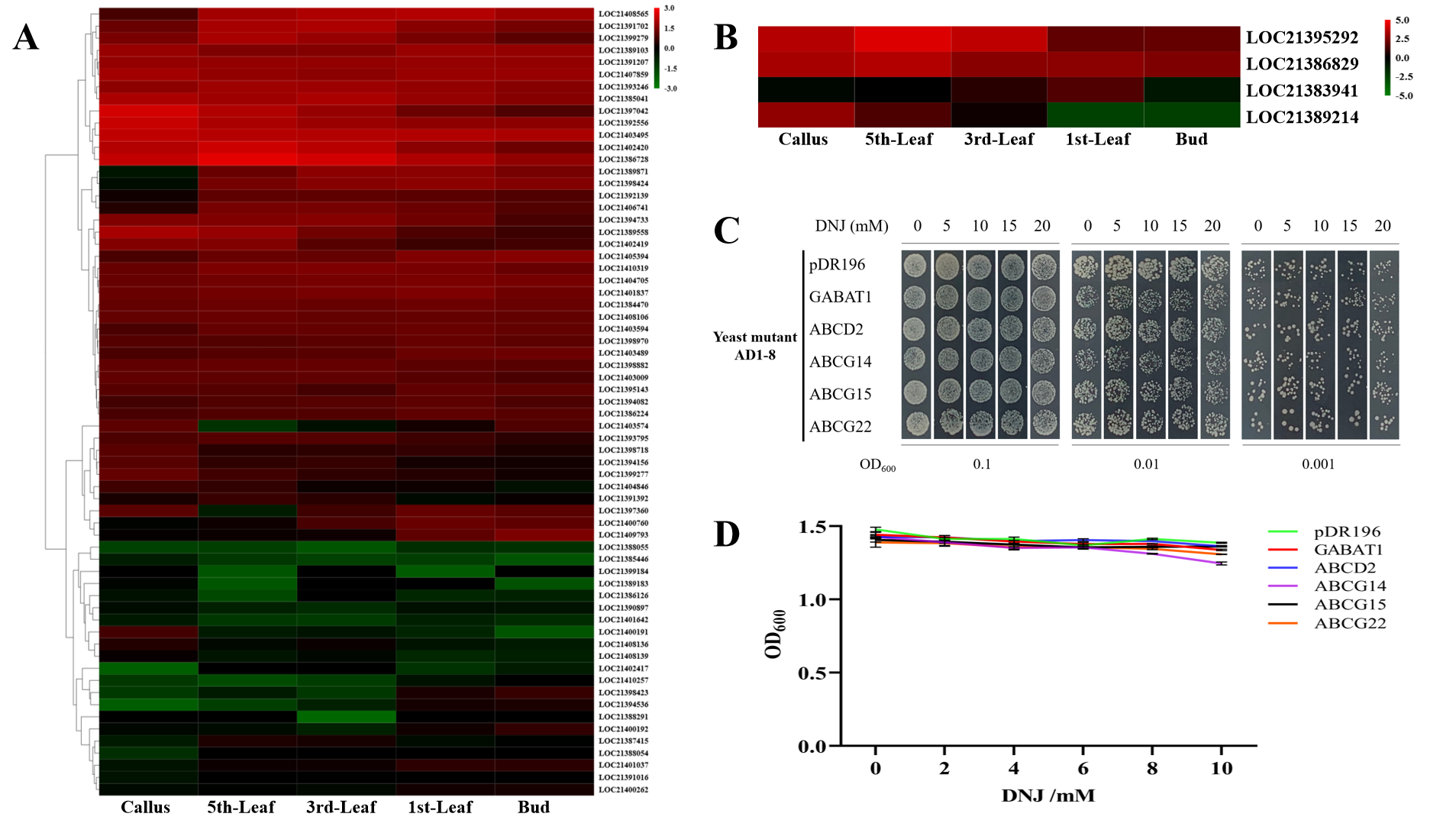

**Figure S4. Tolerance of *S. cerevisiae* strain AD1-8 expressing ABC transporters and GABA transporters at low levels in callus tissue and high levels in leaves in response to DNJ.**

(**A**) Gene expression analysis of ABC transporters in mulberry. Among 65 ABC transporters, the transcripts LOC21406741, LOC21392139, LOC21398424, and LOC21389871, were identified and annotated as ABCD2, ABCG14, ABCG15, and ABCG22, with low expression in callus and high expression in leaves.

(**B**) Gene expression analysis of GABA transporters. One transcript LOC21383941, annotated as GABAT1, exhibited low expression in callus and high expression in leaves.

(**C**) Effect of heterologous expression of GABAT1, ABCD2, ABCG14, ABCG15, and ABCG22 in *S. cerevisiae* strain AD1-8 on tolerance to DNJ.

(**D**) Yeast growth curve for GABAT1, ABCD2, ABCG14, ABCG15, and ABCG22 expressed in liquid medium with the indicated concentrations of DNJ. Data are shown as the mean ± SE (n = 3).

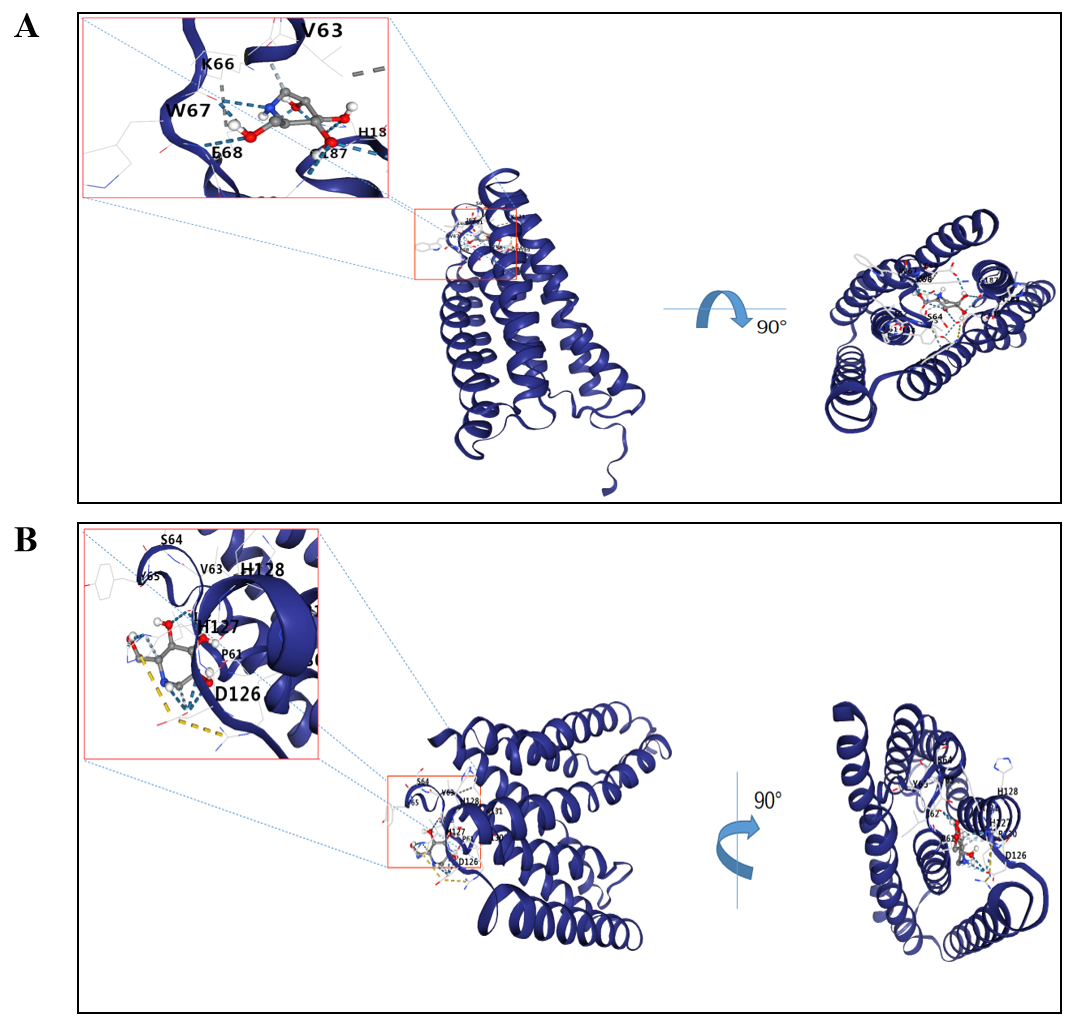

**Figure S5. Predicted structure of SWEET3 docked with DNJ.**

In docking pose A and B, the positively charged imino group of DNJ potentially interacts with (**A**) the 68th residue glutamate and (**B**) the 126th residue aspartate.

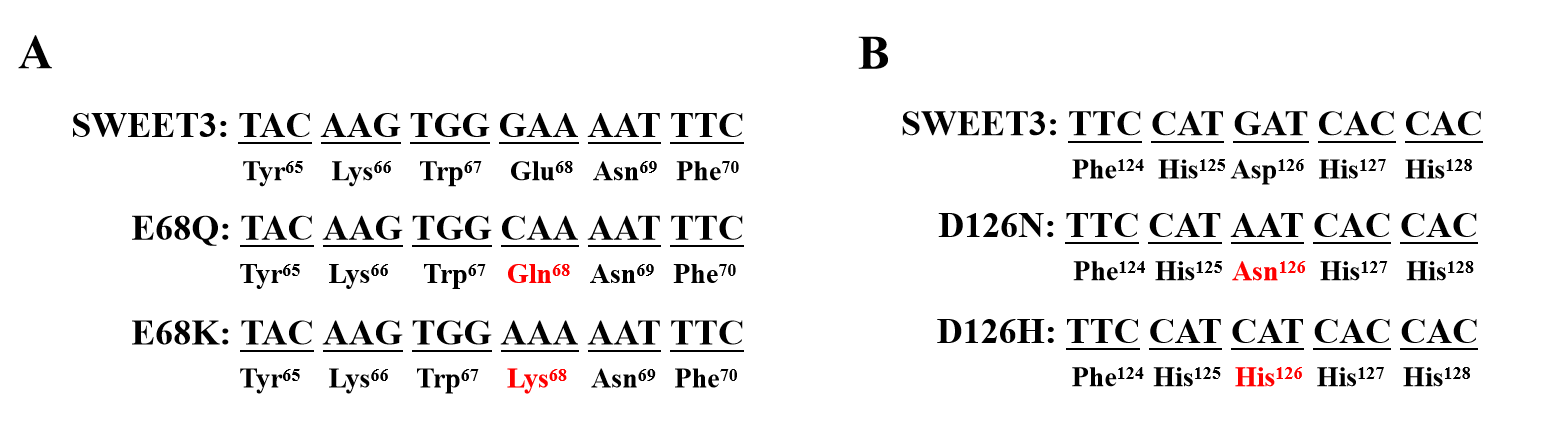

**Figure S6. Sequence alignment of SWEET3, the 68th residue mutants (A), and the 126th residue mutants (B).**

The mutation sites of the sequence are highlighted in red. E68Q, mutation of the 68th Glu (E) to Gln (Q); E68K, mutation of the 68th Glu (E) to Lys (K); D126N, mutation of the 126th Asp (D) to Asn (N); D126H, mutation of the 126th Asp (D) to His (H).

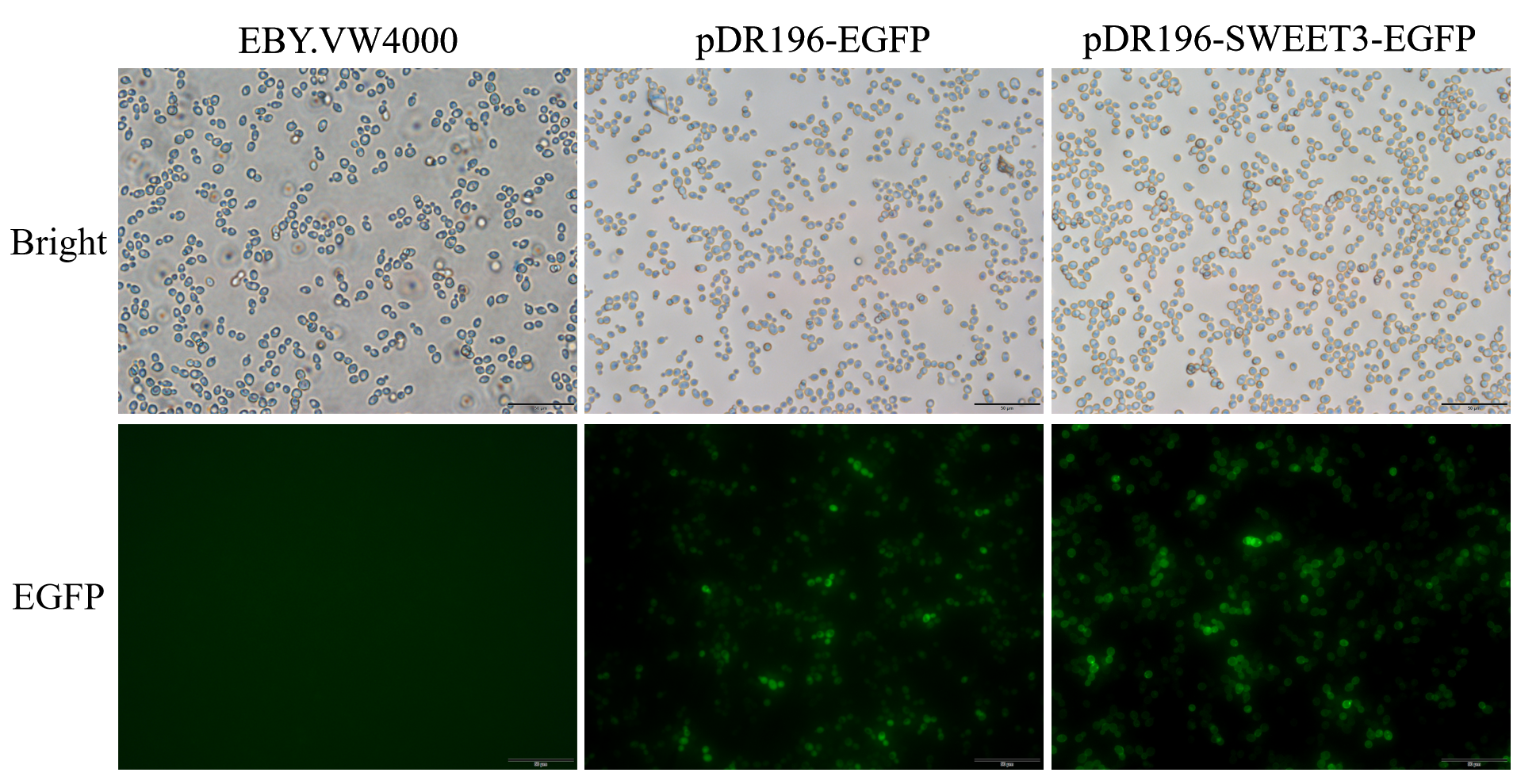

**Figure S7. Subcellular localization of SWEET3 in *S. cerevisiae* strain EBY.VW4000.**

Fluorescence microscope images of *S. cerevisiae* EBY.VW4000 cells expressing pDR196-EGFP or pDR196-SWEET3-GFP. Localization of SWEET3 was captured by fluorescence microscopy, the pDR196 containing the EGFP reporter gene was used as the control, showing the fluorescence signal in the cytoplasm, the SWEET3-EGFP fusion protein exhibited a fluorescence signal localized in the *S. cerevisiae* cell membrane. Bright, bright-feld image. EGFP showed the green fluorescence channel. Scale bars, 50 μm.

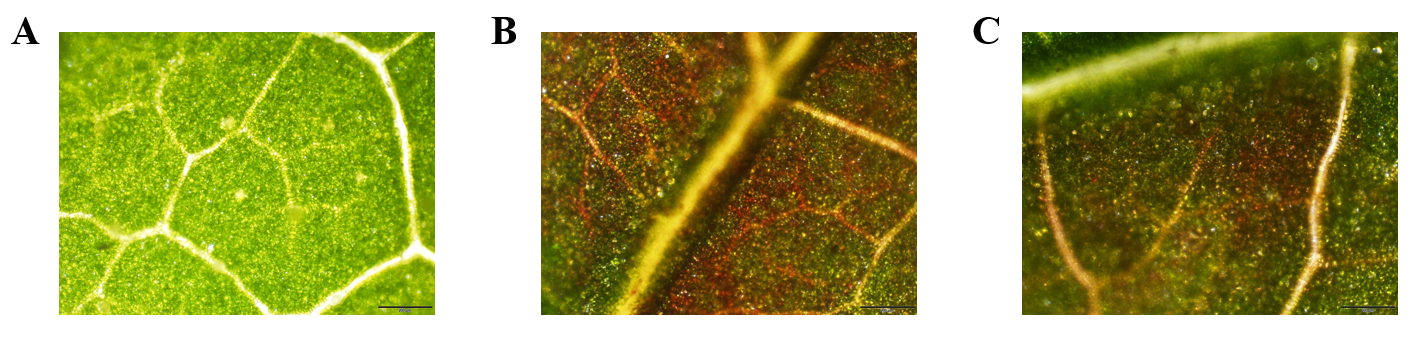

**Figure S8. Transient expression of *RUBY-SWEET3* in mulberry leaves.**

(**A**) Wild type mulberry leaves. Agrobacteria that contain the expression cassette of (**B**) *RUBY* and (**C**) *RUBY-SWEET3* under the control of *CaMV 35S* promoter were infiltrated into mulberry leaves. Scale bars represent 200 μm.
